# A mobile genetic element in the SARS-CoV-2 genome is shared with multiple insect species

**DOI:** 10.1101/2020.06.29.177030

**Authors:** Torstein Tengs, Charles F. Delwiche, Christine Monceyron Jonassen

## Abstract

Unprecedented quantities of sequence data have been generated from the newly emergent severe acute respiratory syndrome coronavirus 2 (SARS-CoV-2), causative agent of COVID-19. We document here the presence of s2m, a highly conserved, mobile genetic element with unknown function, in both the SARS-CoV-2 genome and a large number of insect genomes. Although s2m is not universally present among coronaviruses and appears to undergo horizontal transfer, the high sequence conservation and universal presence of s2m among isolates of SARS-CoV-2 indicate that, when present, the element is essential for viral function.

## INTRODUCTION

Single-stranded RNA viruses, such as coronaviruses, are known to have genomes with strong secondary structural features, such as stem-loop regions and pseudoknots. We have previously reported the presence of a 41-43 nucleotide long hairpin-forming element, referred to as stem-loop II-like motif (s2m) (Jonassen, et al. 1998), in several families of positive-sense single-stranded RNA ((+)ssRNA virus)) viruses (Tengs, et al. 2013). The molecular structure has been mapped in great detail for SARS-CoV (Robertson, et al. 2005). Its sequence and secondary structure are highly conserved, but the phylogenetic distribution among viral genomes is very patchy. These properties indicate that the sequence is mobile, and, to our knowledge, s2m represents the only known example of a genetic element with the ability to move between distantly related viruses. The function and evolutionary origin of s2m remains unknown, but the high level of conservation seen even in distantly related viruses despite relatively high overall mutation rates suggests that s2m is under strong selection. When comparing s2m motifs from different virus species, there are conserved residues both in loop- and base-pairing regions, indicating that there is selective pressure to maintain both the primary and secondary structure. The element is always present near the 3’ end of the genome, and in all virus families where s2m has been reported, there are examples of species carrying two (non-identical) back-to-back copies (Quan, et al. 2010; Tengs, et al. 2013). As noted above, related viruses may lack s2m entirely, but when present, its sequence is always highly conserved.

SARS-CoV-2 (Gorbalenya, et al. 2020) is a member of the SARS-related Sarbecovirus subgenus (Andersen, et al. 2020) and this group of coronaviruses is known to contain s2m (Tengs and Jonassen 2016). The presence of s2m in the SARS-CoV-2 genome (GenBank accession MN908947, position 29727-29768) and other members of this group is probably the result of a single horizontal transfer event, predating the divergence of the SARS-related viruses (Tengs, et al. 2013; Tengs and Jonassen 2016).

## MATERIALS AND METHODS

To characterize the specific genotype of s2m found in SARS-CoV-2, BLASTN (Altschul, et al. 1990) was used to search the entire virus section of GenBank using all s2m sequence genotypes reported in the literature (*n* = 97) (Jonassen, et al. 1998; Robertson, et al. 2005; Tengs, et al. 2013; Tengs and Jonassen 2016) as query sequences. To check for the presence of s2m motifs in insects, both the TSA and the Whole genome shotgun contigs (wgs) databases were mined using the same approach.

For the phylogenetic analyses, sequences were aligned using the Clustal W algorithm (Thompson, et al. 1994) and maximum likelihood analysis were performed using MEGA X (Kumar, et al. 2018). The James-Taylor-Thornton (JTT) substitution model was used with gamma distribution (5 categories) and invariable sites. Branch swapping was done using the subtree-pruning-regrafting method with ‘very strong’ filter.

## RESULTS

A total of 5553 s2m-containing accessions were identified, representing at least four virus families (Supplementary table 1). As expected, there was significant bias towards SARS-CoV-2, with the great majority of the s2m accessions stemming from this species (3984/5553; 72 %). The s2m genotype found in SARS-CoV-2 contains a G > U transversion in position 31 (Figure 1, Figure 2) that is consistent in all available SARS-CoV-2 accessions. This guanine is perfectly conserved outside of the SARS-CoV-2 sequences, with 100 % of the other s2m genotypes having a G in this position (Supplementary table 1, Supplementary table 2).

**Figure 1.**
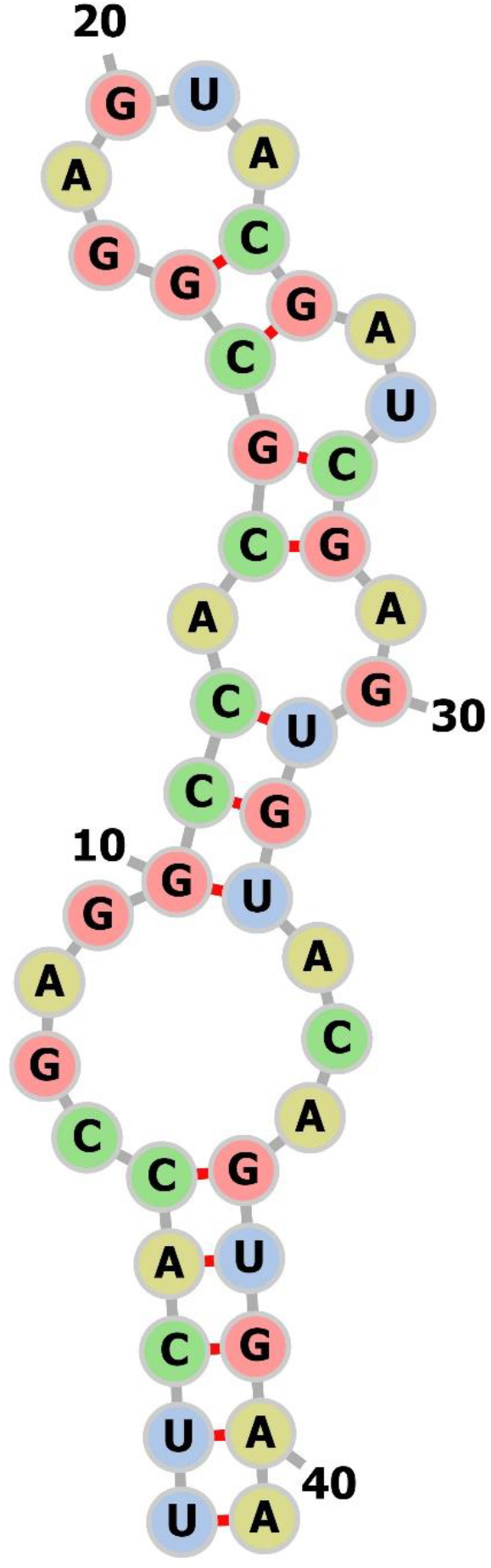
Secondary structure of s2m in SARS-CoV-2 based on SARS-CoV model (Robertson, et al. 2005).

**Figure 2.**
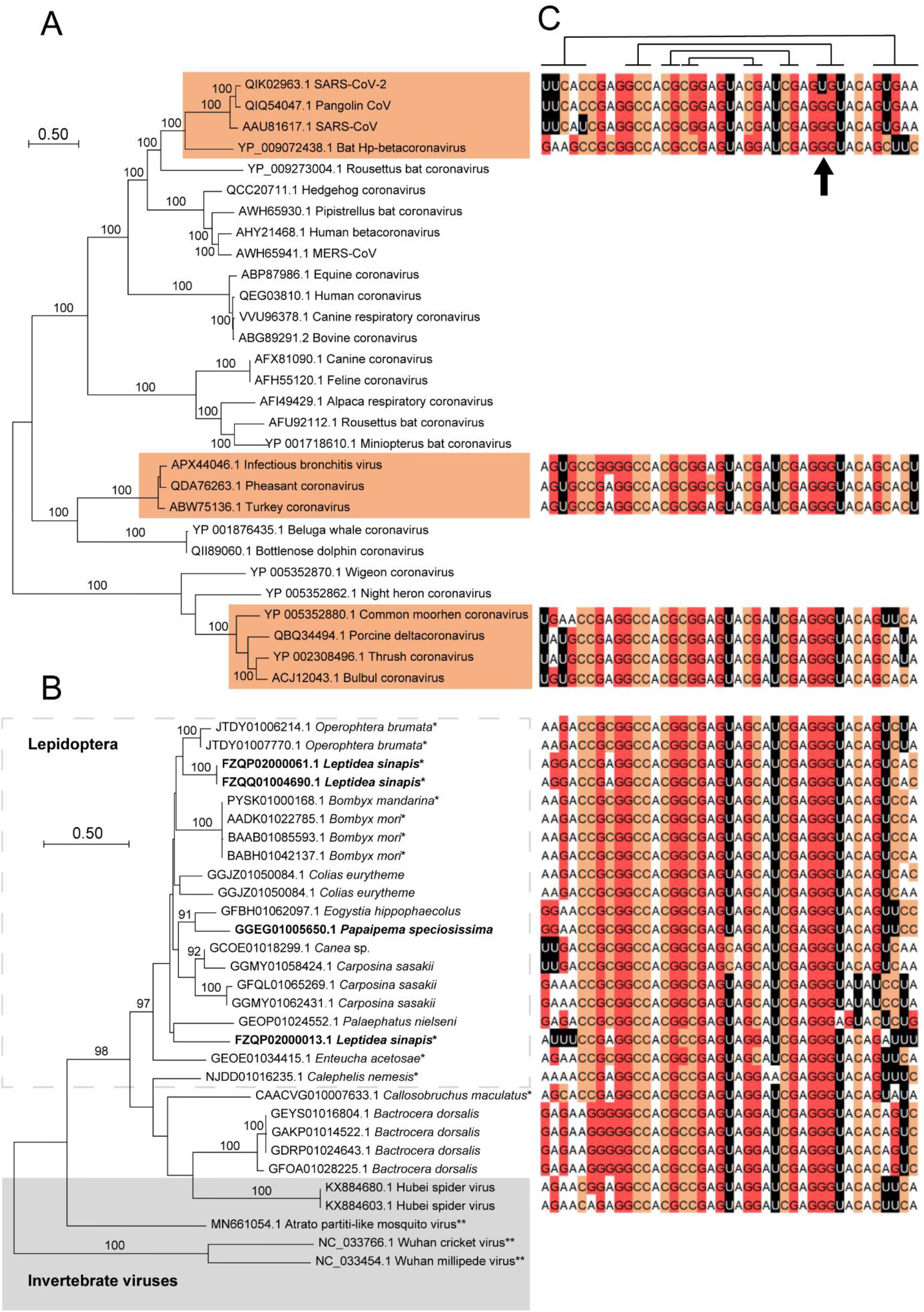
s2m in insects, arachnid viruses and coronaviruses. A) A maximum likelihood analysis was performed on ORF1ab polyprotein sequences from selected coronavirus species. s2m-containing accessions have been highlighted and bootstrap values > 90 % indicated (100 psedoreplicates). B) Maximum likelihood analysis using data from the s2m-associated hypothetical protein (see main text for details). Sequences in boldface stem from reading frames with (multiple) internal stop codons. * - genomic data, ** - accessions without s2m. C) s2m sequences corresponding to operational taxonomic units in the phylogenetic trees. Lines above alignment show (non-canonical) base-pairing residues (Robertson, et al. 2005) and the position with the unique G > U mutation in SARS-CoV-2 has been indicated.

Two of the s2m virus accessions identified were from a recently published RNA-based invertebrate virosphere project (Shi, et al. 2016). These two highly similar sequences, derived from the virome of the spider species *Tetragnatha maxillosa*, encode a hypothetical protein immediately upstream of s2m that could not readily be identified using protein sequence similarity searches. Protein-protein BLAST searches against the non-redundant (nr) GenBank protein database revealed the two best matching non-viral proteins to be from the insect species winter moth (*Operophtera brumata*) and bagworm moth (*Eumeta japonica*). In the *O. brumata* genome sequence, an s2m motif could be identified immediately downstream from the stop codon of a 331 amino acid long uncharacterized protein (Supplementary table 2).

TBLASTN search against the insect section of the GenBank Transcriptome Shotgun Assembly (TSA) database identified a total of 92 protein sequences where > 200 amino acids could reliably be aligned with the *O. brumata* homolog. A phylogenetic analysis of these sequences revealed a complex tree, most likely containing both orthologs and paralogs, but with the *O. brumata* accession and the two spider virus accessions clustering within a well-supported group comprising primarily lepidopteran species (Supplementary Figure 1).

A total of 99 putative s2m-containg insect contigs were identified, representing 45 species (Supplementary table 2). Correlating these findings with the tree topology based on the hypothetical *O. brumata* protein, 18/22 of the insect accessions in the *O. brumata* cluster were found to originate from s2m-carrying insect species (Supplementary Figure 2). None of the accessions outside this cluster showed any traces of s2m. Conversely, proteins encoded by the s2m-containing contigs could reliably be matched with the *O. brumata* protein for 50 (51 %) of the accessions (Supplementary table 2).

A phylogenetic analysis focusing on s2m accessions and using the longest and most similar amino sequences obtained from the TSA, wgs and nr databases (Supplementary table 2) gave a topology that was biased towards lepidopteran species (Figure 2B). Translation of both genomic and transcriptomic data in some cases gave reading frames containing several internal stop codons, but amino acid sequences that could still be aligned to (near) full length (Figure 2B, Supplementary table 2).

## DISCUSSION

As SARS-CoV-2 is embedded within the Sarbecoviruses, it is likely that the unique G > U mutation has occurred specifically during the evolution of the current pandemic strain of SARS-CoV (Figure 2C). An Australian isolate with a 10 base deletion in s2m was also discovered (accession MT007544). Intriguingly, the deletion occurred after passaging isolates in Vero cells (Caly, et al. 2020). This could represent an attenuated SARS-CoV-2 strain, as culturing in permissive cell lines may have altered the selective pressure and alleviated the need to maintain a functional version of the motif.

In the four virus families where s2m has previously been described, the flanking protein is easily recognizable. The protein found in insect species, as well as the two *T*. *maxillosa* viruses, does not appear to have homologs in any of the better-characterized virus groups, nor does it appear to contain any recognizable motifs. A protein secondary structure homology search (Drozdetskiy, et al. 2015) indicated that the *O. brumata* protein may contain an integrase catalytic domain, which would be consistent with the mobile nature of s2m. We were unable to identify any signature of a retroviral origin of the s2m loci in insect genomes, or any other indication of the insect s2m contigs being of viral origin. In several of the genomic contigs, the open reading frame (ORF) covering the uncharacterized protein also appeared to contain introns, generally considered a hallmark of eukaryotic genes. In addition, PCR and Sanger sequencing was performed to confirm the presence of s2m and the upstream ORF in the *O. brumata* genome (Supplementary figure 2), making it very unlikely that the downloaded sequence data stem from (RNA) viruses and do not represent *bona fide* insect sequences.

The insect species that contain s2m (and the associated protein) are distantly related, indicating either a deep evolutionary origin with multiple losses or that this genetic construct is also a mobile element, perhaps using viruses as a vector (Gilbert and Cordaux 2017). The *T. maxillosa* virus could represent such a vector, albeit no s2m sequences or proteins similar to the *O. brumata* protein could be found in any arachnid species using sequence similarity searches. Mobility of genetic elements such as transposable elements (TEs) has previously been reported in insects (Peccoud, et al. 2017), and an analysis of the genomic s2m contigs using the Dfam portal (Hubley, et al. 2016) revealed that several of the accessions had regions with a significant degree of similarity with the long interspersed nuclear element (LINE) L2-1_Ldor_D, previously reported from butterflies (Ray, et al. 2019). The exact evolutionary link between the xenologs of s2m and the unknown protein found in insects and viruses can not be established based on our data. The s2m genotypes found in insects and viruses have similar primary sequence profiles, albeit there appear to be some subtle differences (Figure 3).

**Figure 3.**
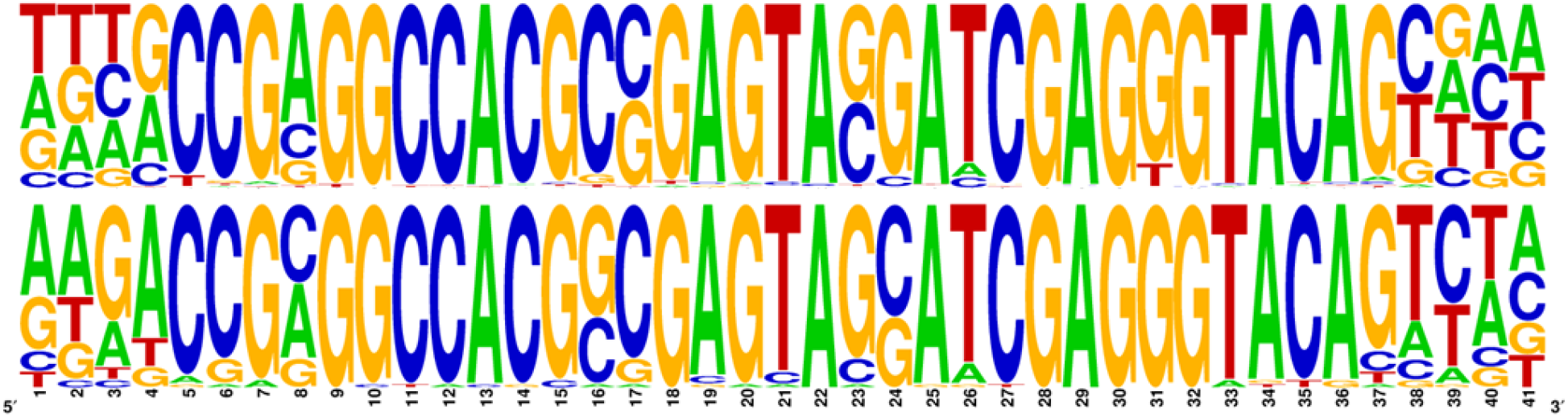
DNA logo frequency plot for s2m genotypes found in viruses (top panel) and insects (bottom panel). Generated using WebLogo (Crooks, et al. 2004).

We believe that the most likely mode of transfer for s2m in viruses is through non-homologous recombination between RNA molecules. Outside the astroviruses, SARS-CoV and SARS-CoV-2 represent the only known examples of s2m-carrying viruses that infect humans, but it seems probable that s2m is still evolutionary active and that this element will continue to affect the evolution of (+)ssRNA viruses. Because the clade of s2m-containing coronaviruses that includes SARS-CoV and SARS-Cov-2 also includes viruses isolated from bat and pangolin, it is unlikely that s2m has played a direct role in zoonotic transfer, but its high degree of sequence conservation suggests that it may present a target for therapy, and the presence of closely related systems in insect hosts provides an opportunity to study the biology of this apparent mobile element in relatively tractable experimental systems.

## ACKNOWLEDGEMENTS

The authors would like to thank M.Sc Anbjørg Rangberg (Østfold Hospital Trust) for help with PCR and sequencing and Dr. Snorre Hagen (Norwegian Institute of Bioeconomy Research) for providing the *Operophtera brumata* DNA. This work was supported by the University of Maryland Agricultural Experiment Station (CFD).

---

**Supplementary figure 1.**
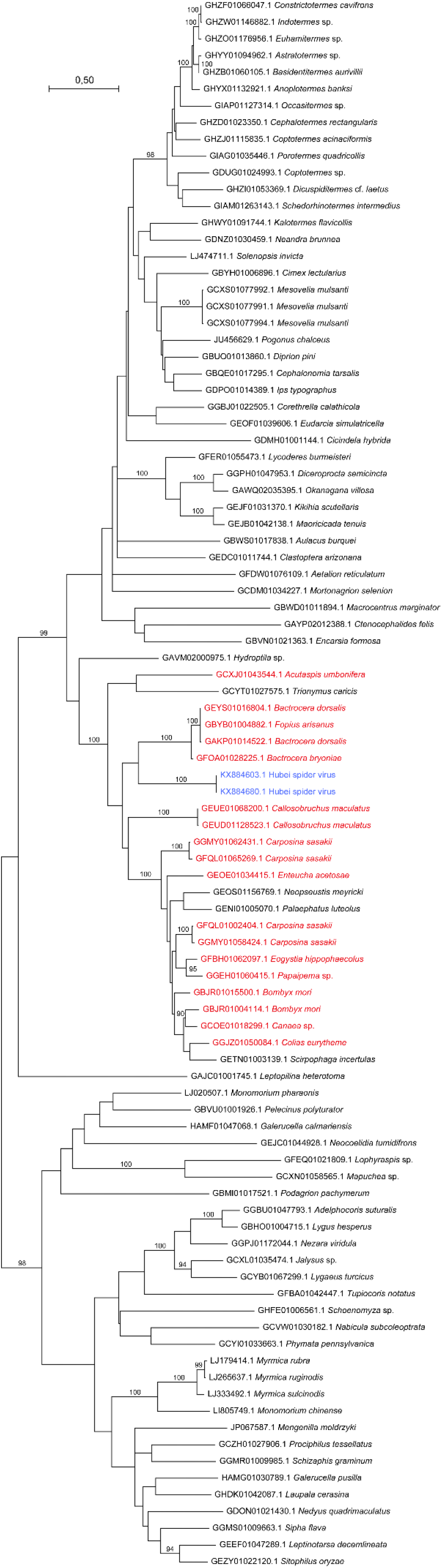
Phylogenetic tree based the hypothetical protein found in *Operophtera brumata* and related insect species. Identical sequences from the same species were removed and a maximum likelihood analysis was performed using MEGA X (Kumar, et al. 2018) after aligning amino acid sequences using the Clustal W algorithm (Thompson, et al. 1994). The James-Taylor-Thornton (JTT) substitution model was used with gamma distribution (5 categories) and invariable sites. Branch swapping was done using the subtree-pruning-regrafting method with ‘very strong’ filter. s2m-containing accessions have been indicated (red: insect species, blue: invertebrate viruses) and bootstrap values > 90 % are shown (100 psedoreplicates).

**Supplementary figure 2.**
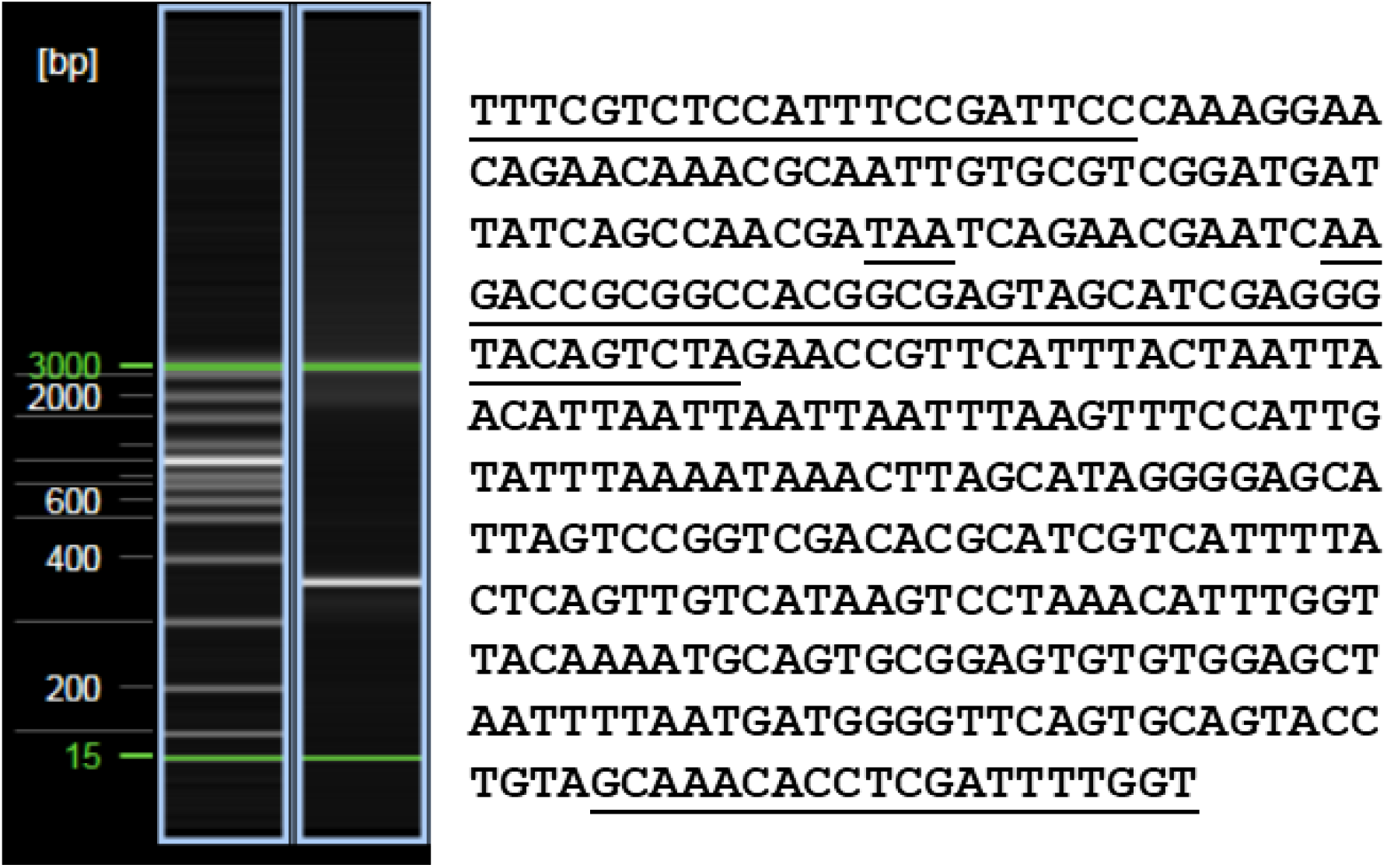
PCR amplification and Sanger sequencing of the s2m locus in the *Operophtera brumata* genome. PCR was performed using the AmpliTaq Gold 360 PCR Master Mix (Thermo Fisher Scientific, Oslo, Norway). Primer sequences, ORF stop codon (TAA) and s2m sequence have been underlined.

